# Mapping the global distribution and spread of the *Plasmodium vivax*-associated virus MaRNAV-1

**DOI:** 10.64898/2026.02.26.708358

**Authors:** Mary E. Petrone, Justine Charon, Rhys H. Parry, Matthew J. Grigg, Kim A. Piera, Jacob A. F. Westaway, Kayoko Shioda, Bruce Russell, Ric N. Price, Timothy William, Enny Kenangalem, James S McCarthy, Bridget E. Barber, Edward C. Holmes, Nicholas M. Anstey

## Abstract

Matryoshka RNA virus 1 is a bi-segmented and single-stranded RNA virus associated with *Plasmodium vivax*, a cause of human malaria. Little has been uncovered about the epidemiology and ecology of this virus since its discovery in 2019. To address this, we used a combination of primary and publicly available metatranscriptomic data to map the geographic distribution and host associations of MaRNAV-1. We detected this virus throughout Southeast Asia, in parts of South America, and, for the first time, in Oceania. Despite its broad distribution, MaRNAV-1 was found exclusively in metatranscriptomes containing *P. vivax*, suggesting that there is a specific virus-host relationship that has shaped the evolutionary history of this virus. We were unable to estimate the emergence date of the MaRNAV-1 lineage; however, phylogeographic mapping analysis suggested that MaRNAV-1 may have radiated from Southeast Asia. Our findings have both evolutionary and public health implications and can serve as the basis for future investigations in these fields.

## INTRODUCTION

The first RNA virus to be associated with *Plasmodium*, a single-celled parasite that can cause malaria in humans, was discovered in 2019^1^. Matryoshka RNA virus 1 (MaRNAV-1) is a bi-segmented and single-stranded RNA virus from the family *Narnaviridae* (phylum *Lenarviricota*) that, to date, is exclusively associated with the species *P. vivax*. MaRNAV-1 has been identified in *P. vivax*-infected human and mosquito blood samples from Malaysia, Cambodia, Thailand, and Colombia^1^. Since its discovery, six additional lineages of MaRNAVs (MaRNAV-2-7) have been identified^1-4^. Collectively, these exhibit broader diversity and host range than MaRNAV-1. Identified in birds sampled in Oceania, North America, and Europe, MaRNAVs 2-7 are presumed to infect species of *Leucocytozoon* and *Haemoproteus sp*. which, along with *Plasmodium*, are members of the order Haemosporida (phylum Apicomplexa).

That only one MaRNAV species has been identified in *Plasmodium* compared to four in leucocytozoa and two in Haemoproteus indicates an important gap in our knowledge of virus evolution. The identification of MaRNAVs in multiple genera of Haemosporida suggests that this viral lineage was present in the common ancestor of the order, which is estimated to have originated ∼70 million years ago around the time of the Cretaceous–Paleogene (K–Pg) mass extinction event^5^. However, Charon, *et al*., found no evidence of MaRNAVs in the *P. knowlesi* and *P. falciparum* parasites that were co-circulating with the MaRNAV-positive *P. vivax* infected-samples in their cohort^1^. One explanation is that this was the consequence of the small sample size (n=18) and limited catchment area (Sabah, east Malaysia) of this initial study. Alternatively, the life cycle of *P. vivax*, which involves the establishment of a latent infection in the liver that can reactive after initial infection and cause relapsing episodes of malaria and ongoing transmission of the parasite, may facilitate infection by MaRNAV-1 or have its relapse frequency modulated by viral infection^4^. *P. ovale* and *P. cynomolgi* are the only other human-infecting species that exhibit a latent hepatic hypnozoite stage^6^. Hence, such a feature might render *P. vivax* susceptible to MaRNAV infection or less equipped to mount a protective response against it compared to other *Plasmodium* species.

The changing ecology of *P. vivax* might also impact the distribution and prevalence of MaRNAV-1. Although *P. falciparum* accounts for >90% of malaria cases globally^7^, successful elimination strategies in co-endemic areas are leading to a relative increase in the proportion of malaria due to *P. vivax*. Indeed, *P. vivax* has become the predominant cause of malaria in Southeast Aisa, the Horn of Africa, and the Americas^8^. *P. vivax* is more difficult to eliminate than *P. falciparum*, since it requires radical cure of all stages of the parasite. These challenges are compounded by the emergence and spread of chloroquine resistance and the need for prolonged treatment course of primaquine to kill liver stages^9-13^. Whether the shift in the epidemiology of *P. vivax* corresponds to an increase in MaRNAV-1 has not been assessed, in part because no baseline measurements have been recorded. Specifically, the frequency at which MaRNAV-1 co-occurs with *P. vivax* is not known, nor if this frequency varies by geographic region.

Herein, we aim to help fill this gap by extending our search for MaRNAV-1 to three additional species of human-infecting *Plasmodium* species collected across southeast Asia, Africa, South America, and Oceania. In doing so, we consider three hypotheses that could explain the evolutionary history and phylogenetic distribution of MaRNAV-1: (1) MaRNAV-1 can infect all *Plasmodium* species, and this will be revealed by screening additional species from a broader geographic range; (2) MaRNAV-1 emerged in *P. vivax* after this species diverged from other members of its genus; (3) MaRNAVs were present in the *Plasmodium* common ancestor, but non-vivax *Plasmodium* species evolved mechanisms to prevent infection by MaRNAV-1. We leverage metatranscriptomic sequencing and time-scaled phylogenetics to investigate the geographic and putative host range of MaRNAV-1, thereby evaluating the plausibility of each hypothesis.

## RESULTS

### The distribution of MaRNAV-1 in *P. vivax* is global and varied

To establish a reference for the geographic range and frequency of MaRNAV-1, we began by measuring these metrics in *P. vivax* metatranscriptomes. Accordingly, we compiled a data set of *P. vivax* metatranscriptomes from (i) primary samples (n=34, including 7 that were previously described by Charon *et al*.^1^; **Supp. Table 1**) and (ii) publicly available RNA libraries from the National Center for Biotechnology Information (NCBI) Sequence Read Archive (SRA) (n=1,431, as of May 2025; **Supp. Table 2**). Screening these libraries for MaRNAV-like viruses using DIAMOND BLAST^14^ returned 610 putative MaRNAV RNA-dependent RNA polymerase (RdRp) sequences (**Supp. Data 1**) with minimal sequence diversity (at least 95% amino acid similarity to the original MaRNAV-1 RNA-dependent RNA polymerase (RdRp) [acc. QGV56801] and 97% amino acid similarity to the original MaRNAV-1 hypothetical protein [acc. QGV56796]). Relatives of MaRNAVs-2-7 were not detected. We classified all putative viruses identified as MaRNAV-1.

Our search expanded the known geographic and temporal distribution of MaRNAV-1, revealing the first instances of MaRNAV-1 in *Plasmodium* libraries from Peru (n=27 of 319), Indonesia (n=14 of 25), Solomon Islands (n=1 of 2), and Papua New Guinea (n=1 of 3) (**Fig. 1**). The presence of this virus in human blood isolates from Peru suggested that virus-infected *Plasmodium* have been transmitted to humans by mosquito vectors in South America. Previously, MaRNAV-1 virus had only been detected in mosquitoes collected in Colombia^1^. We also detected MaRNAV-1 in the earliest libraries in our data set (Indonesia, 2004), extending its known circulation in Southeast Asia by 10 years. Fragments of the hypothetical protein were recovered in two of the 27 libraries we screened from Brazil without corresponding RdRp segments. We did not consider this as sufficient evidence that MaRNAV-1 is circulating in the country, but additional sampling could resolve this given that the virus is present in neighbouring Colombia and Peru. We did not identify MaRNAV-1 in the one *P. vivax* library available from Africa (Ethiopia).

**Figure 1.**
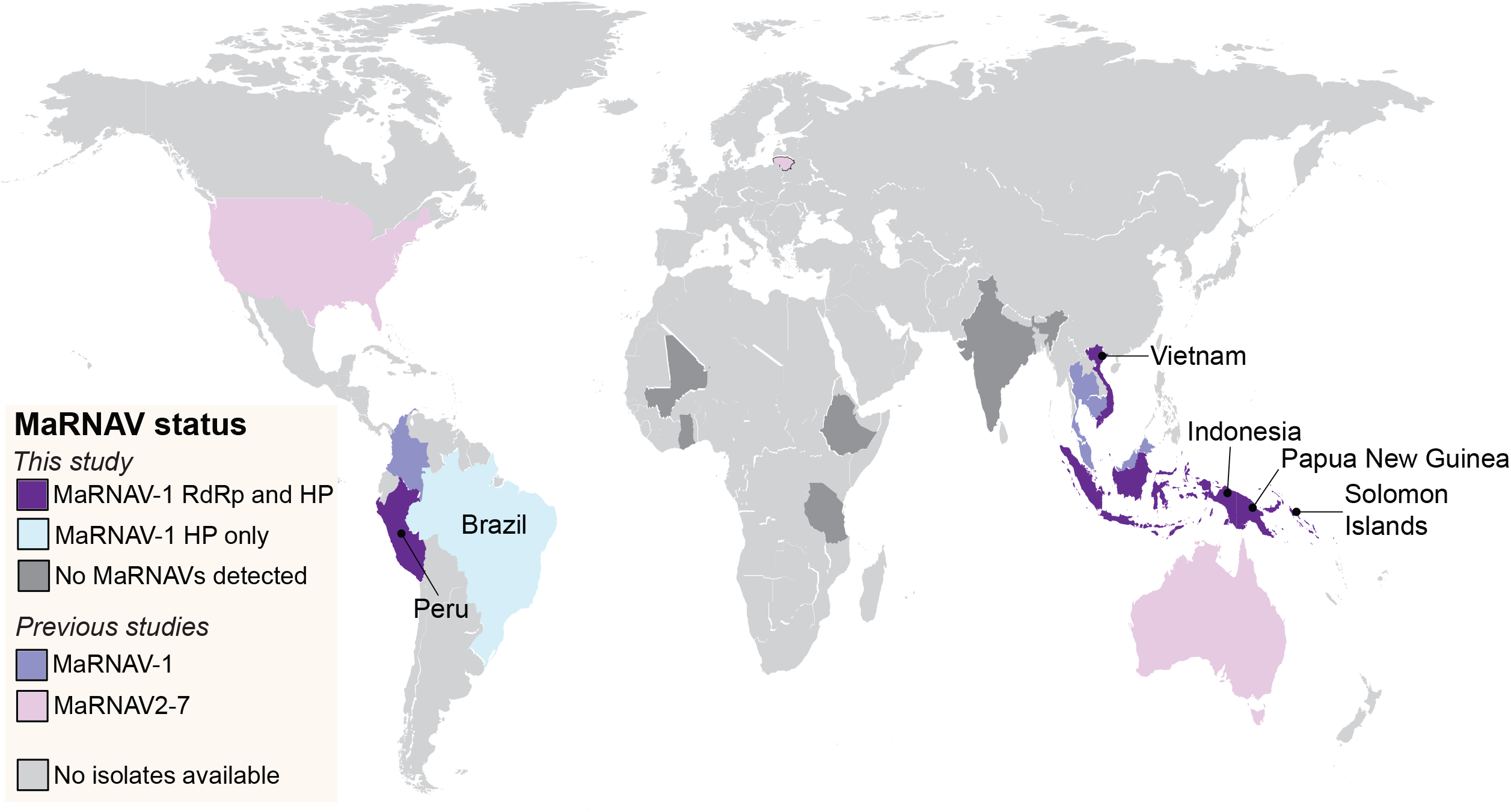
Global distribution of MaRNAVs in Haemosporida transcriptomes. MaRNAVs 2-7 were previously detected in USA^4^, Australia^1^, and Lithuania^2^ (pink). MaRNAV-1 was previously detected in Colombia, Malaysia, and Cambodia by Charon, *et al*.^1^ Abbreviations: RdRp (RNA-dependent RNA polymerase), HP (hypothetical protein)

The frequency of MaRNAV-1 in *P. vivax* libraries varied considerably. Among the primary isolates, frequency ranged from 0% (India, 0 out of 1) to 77.8% (Papua, Indonesia, 14 out of 18) (**Table 1**).

**Table 1:** Frequency of MaRNAV-1 in primary human blood isolates infected with *P. vivax*.

| Country | % positive | # positive samples | # total samples |
| --- | --- | --- | --- |
| India | 0 | 0 | 1 |
| Indonesia | 77.8 | 14 | 18 |
| Malaysia | 61.5 | 8 | 13 |
| Solomon Islands | 50 | 1 | 2 |

The frequency of MaRNAV-1 in libraries available on the SRA was more difficult to estimate because they were derived from a combination of human and mosquito samples and we could not assume that individual libraries represented individual infections due to pooling. Therefore, frequency estimates from the SRA could not be directly compared to our findings of the primary samples. Despite this, we again observed varying frequencies of MaRNAV-1 in clinical samples from individual countries, albeit at a lower rate than what we observed in the primary samples (**Table 2, Supp. Table 2**). Overall, our observations suggest that MaRNAV-1 is present in human-infecting *P. vivax* on at least three continents.

**Table 2:**
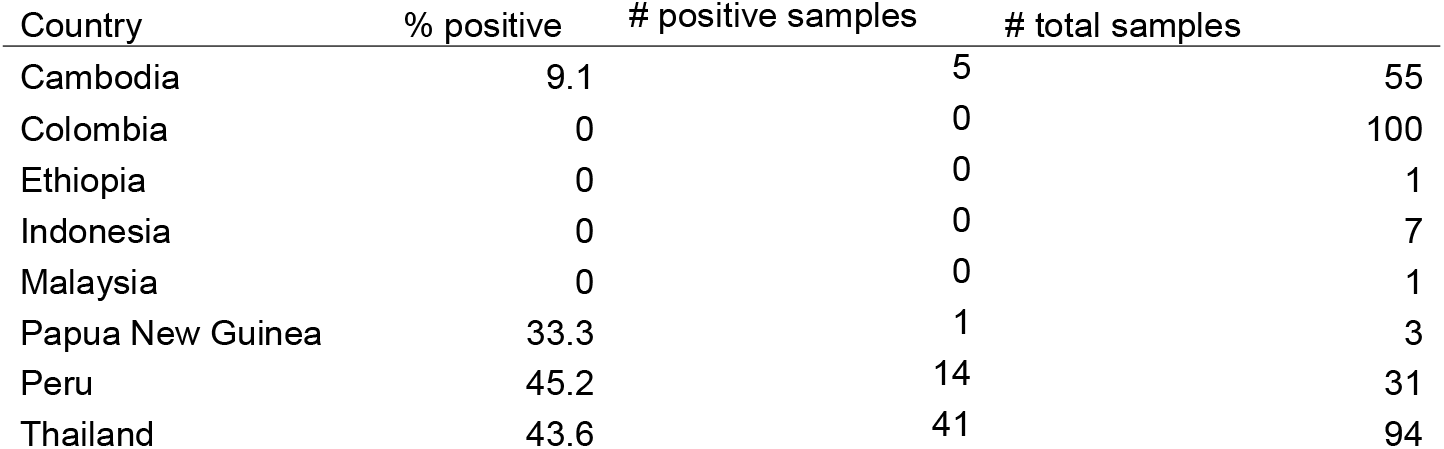
Frequency of MaRNAV-1 in publicly available *P. vivax* transcriptomes generated from clinical isolates of human blood.

### MaRNAV-1 is specific to *P. vivax*

Having previously shown an absence of MaRNAV-1 in small numbers of other *Plasmodium* species in Sabah, Malaysia (*P. falciparum* and *P. knowlesi*)^1^, we next assessed whether MaRNAV-1 is similarly absent in a broader range of non-vivax *Plasmodium* species, including the relapsing species *P. ovale* and *P. cynomolgi*. We tested this by screening primary samples from patients who had non-*P. vivax* malaria from the same catchment area in Southeast Asia in which MaRNAV-1 was first detected^1^ and in patients from the Pacific and Africa (**Fig. 1**). These samples comprised 13 human blood samples with documented *P. falciparum* from Sabah, Malaysia (n=5), Papua, Indonesia (imported; n=2), Cambodia (n=1), Mali (n=1), Ghana (n=1) and Tanzania (n=2); 8 samples with documented *P. malariae* from Sabah (n=2) and Papua, Indonesia (n=6); and 7 samples with documented *P. ovale* infections from Papua, Indonesia (n=6) and Sabah (n=1) (**Supp. Table 1**). We also screened a laboratory-adapted strain of *P. cynomolgi* Berok^15^. Finally, we screened all publicly available *Plasmodium* metatranscriptomes that were not previously captured by Charon, *et al*^1^., (n=7,662 available on NCBI as of October 2025; **Supp. Tables 3 [*P. falciparum*; n = 5**,**753], 4 [non-*P. falciparum*; n = 1**,**909]**). These included transcriptomes of laboratory strains (e.g., *P. falciparum* 3D7) and laboratory-culture mouse strains (e.g., *P. berghei*) in addition to natural human infections.

This analysis did not identify MaRNAV-1 in any of the non-*P. vivax* transcriptomes of the primary clinical isolates, including those that were co-circulating with infected *P. vivax* isolates. Specifically, we found no evidence of MaRNAV-1 in *P. malariae, P. ovale*, or *P. falciparum* parasites that were co-circulating with *P. vivax* in Papua, Indonesia between 2004 and 2016 (including imported cases), a period during which we repeatedly detected MaRNAV-1 in *P. vivax* infections (**Fig. 2a**). In addition, we did not identify MaRNAV-1 in any of the seven isolates from patients infected with *P. ovale*, which can establish latent liver stages and cause relapsing malaria.

**Figure 2.**
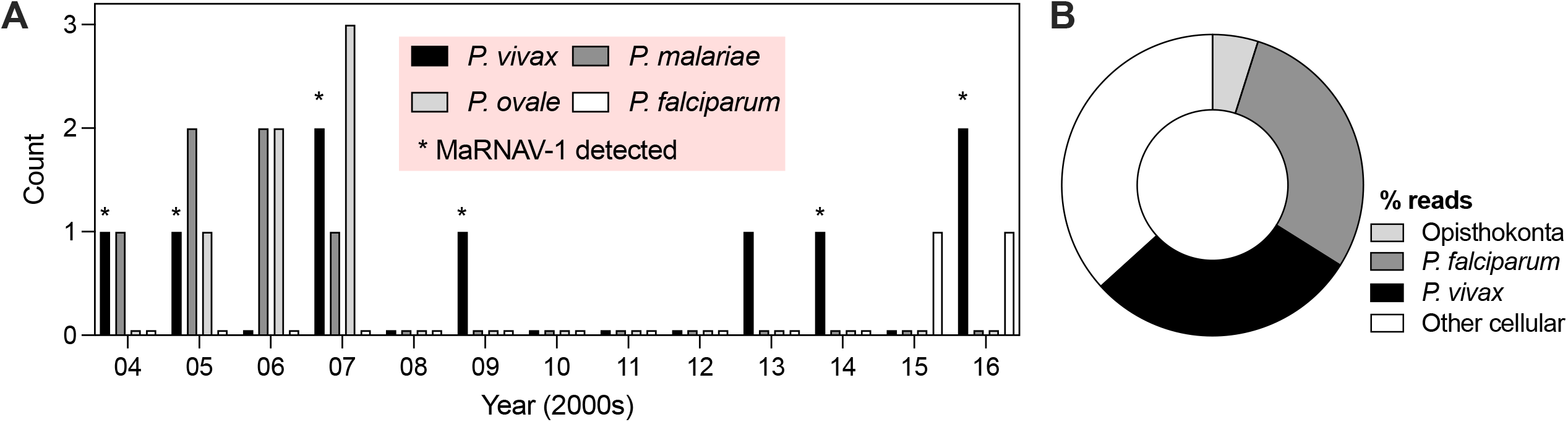
Evidence for MaRNAV-1 specificity to *P. vivax*. (a) Number of primary human-infecting *Plasmodium* species collected per year originating in Papua, Indonesia (including imported cases). The MaRNAV-1 RdRp segment was only detected in *P. vivax* samples (asterisks); (b) Representative output of Kraken2 analysis showing the composition of MaRNAV-1-positive *P. falciparum* library (SRA ID: SRR25922176).

Interestingly, however, we detected the MaRNAV-1 RdRp in 120 *P. falciparum* libraries (8.3%), including 9 collected in Vietnam (**Supp. Table 3**). To assess whether their presence could be *bona fide* evidence of MaRNAV-1 association with *P. falciparum*, we first considered whether these libraries were co-infected with *P. falciparum* and *P. vivax*. This was plausible because all MaRNAV-1-positive *P. falciparum* libraries were generated as part of studies for which co-infection was not an exclusion criterion (BioProjects PRJNA716963, PRJNA1013216, and PRJDB2573)^16,17^, and all three studies were carried out in Southeast Asia, where both *P. falciparum* and *P. vivax* co-circulate^18^. Additionally, no MaRNAV-1 reads were detected in libraries from samples from the African continent (where *P. vivax* transmission is low and spatially heterogeneous) or in laboratory-adapted strains, which would be known to not contain mixed species. To test for co-infection, we screened the 120 *P. falciparum* libraries containing MaRNAV-1 RdRp sequences for reads aligning to the *P. viva*x genome (**Supp. Data 2**). We found evidence of *P. vivax* in all 120 *P. falciparum* libraries, including 23 instances in which *P. vivax* was equal to or more abundant in the library compared to *P. falciparum* (e.g., **Fig. 2b**). We also calculated the probability of finding MaRNAV-1 in 8.3% or less of human-infecting *P. falciparum* if the true prevalence was consistent with that of MaRNAVs in publicly available *P. vivax* transcriptomes (weighted mean=20.9%, **Table 2**). To do this, we applied a binomial distribution function (***Eq. 1***), which returned a one-sided p-value of <0.0001: the probability of observing 120 or fewer positives in 1453 samples if the true prevalence was 20.89% was effectively zero. From these two lines of evidence, we concluded that only one MaRNAV species (MaRNAV-1) is associated with human-infecting *Plasmodium*, and it is specific to *P. vivax*.

### Southeast Asia as the driver of MaRNAV-1 spread

We next sought to determine when and from where MaRNAV-1 emerged in *P. vivax*. A recent emergence date would be consistent with the hypothesis that MaRNAV-1 evolved after the divergence of *P. vivax* from a common *Plasmodium* ancestor.

We began by placing the phylogenetic diversity of the novel MaRNAV-1 viruses discovered in this study within all previously identified MaRNAVs and the 100 most closely related RdRp sequences to the MaRNAV-1 RdRp. This analysis placed MaRNAV-4 and MaRNAV-5 in a separate lineage with non-MaRNAV members of the *Narnaviridae* (**Fig. S2**). Consistent with Esperanza, *et al*., who could not detect a second segment in the corresponding libraries^4^, we concluded that these viruses should not be classified as MaRNAVs and excluded them from further analysis of this group. However, this does not rule out their association with *Haemoproteus*.

To help determine from where MaRNAV-1 may have emerged, we performed a phylogeographic mapping analysis. Viruses from Southeast Asia were polyphyletic with no defined country-specific clustering (**Fig. 3**). Although some country-specific clades were present, particularly for Thailand, sequences from Cambodia and Vietnam were dispersed throughout the phylogeny, likely a reflection of their close geographic relationships. Similarly, viruses from Oceania (Solomon Islands and Papua New Guinea) fell in different parts of the phylogeny, yet among viruses from Southeast Asia, consistent with separate introductions from this region. In contrast, viruses from South America formed a monophyletic group among Southeast Asian viruses, indicative of a single introduction (**Fig. 3**). This analysis also supports the conclusion that the MaRNAV-1 sequences detected with putative *P. falciparum* libraries were associated with *P. vivax* because MaRNAV-1 from both host species were distributed throughout the MaRNAV-1 clade rather than forming distinct lineages as has been observed in leucocytozoan-associated MaRNAVs (**Fig. S2**).

**Figure 3.**
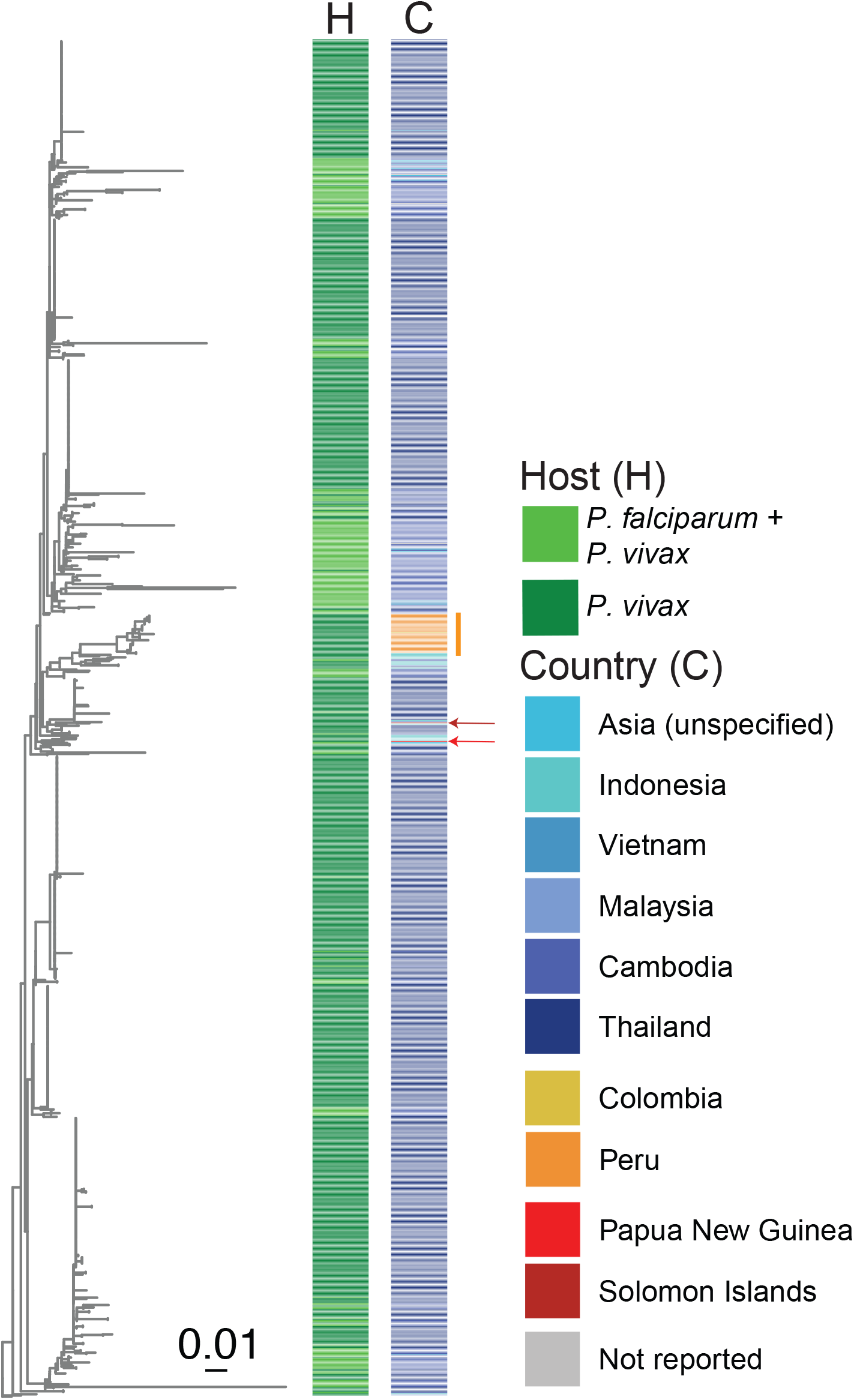
Phylogenteic distribution of MaRNAV-1 by country. Midpoint rooted maximum likelihood phylogenetic tree of the RdRp rooted on the Leucocytozoon-associated MaRNAV lineages. Arrows indicate placement of MaRNAV-1 RdRp sequences identified in Oceania. The vertical orange bar corresponds to samples collected in South America. Branches are scaled according to number of amino acid substitutions.

We were similarly unable to estimate the precise date of emergence of MaRNAV-1 relative to the divergence of *P. vivax*. An analysis of root-to-tip genetic distances against year of sampling using a refined data set of tips with known collection dates did not return a detectable clock-like structure (R-squared = 2.55E-2 with heuristic residual mean squared function for the best-fitting root; **Fig. S3**), likely reflecting the short sampling period (∼15 years). Sampling of *Plasmodium* collected prior to the 21^st^ century may be needed to resolve this limitation.

## DISCUSSION

Our study presents strong evidence for a specific association between MaRNAV-1 and *P. vivax*. Our data also suggest that MaRNAVs are not associated with other human-infecting *Plasmodium* species, including *P. falciparum* and relapsing species such as *P. ovale*, suggesting that MaRNAV-1 is unable to infect all *Plasmodium* species. This apparent specificity contrasts with the four distinct MaRNAV lineages that have been reported in leucocytozoans and two related viruses in *Haemoproteus*. Our *P. vivax* metatranscriptome data set contained a disproportionate number of samples from Southeast Asia, but we were still able to conclude that this region is the likely driver of global spread of MaRNAV-1-infected *P. vivax*, as evidenced by the monophyly of viruses from South America and the two introductions of MaRNAV into Oceania from Southeast Asia (**Fig. 3**).

More granular assessment of historical MaRNAV-1 circulation patterns will require robust historical sampling and better representation of the African continent. The earliest samples in our data set were from 2004, and this limited time span was insufficient to allow the emergence of a temporal signal in our root-to-tip analysis. We could not determine whether the arrival of MaRNAV-1 in South America coincided with the first introduction of *P. vivax* to the continent (the timing of which is still debated^19^). Similarly, whether MaRNAV-1 is present on the African continent could not be ascertained because only one *P. vivax* library from the continent was available for our analysis (Ethiopia). A broader sampling is therefore necessary to determine when MaRNAV-1 emerged in *P. vivax* relative to the spread of this cellular pathogen around the world.

Despite these limitations, the heterogeneity of MaRNAV-1 status with *P. vivax* populations suggests that MaRNAV-1 may be transmissible between *P. vivax* hosts, which would be a unique feature amongst the *Narnaviridae*. Canonical members of the *Narnaviridae* were so named because their genomes comprise a single gene encoding the RdRp (hence the name “naked RNA” – narna). The lack of capsid is likely to limit the possibility of horizontal transmission. Indeed, it has been hypothesised that capsid-less RNA viruses are transmitted vertically, persisting exclusively in the cytoplasm of the cells they infect^20,21^. Interestingly, however, MaRNAVs are bi-segmented, encoding a ‘hypothetical protein’ that could have a structural purpose. If the hypothetical protein is indeed a capsid-like protein that facilitates the transmission of MaRNAVs between protozoan hosts, its acquisition would represent a significant event in the evolution of the MaRNAV viral lineage, possibly even marking its emergence. Identifying the source of this protein and when it was acquired would generate insights into the evolution of the *Narnaviridae* and potentially virus-protozoa interactions if the second segment was derived from a host gene. It could also mark an evolutionary commitment to a specific host type, just as the acquisition of the E1/E2 glycoprotein in some members of the *Flaviviridae* appears to be linked to commitment to vertebrate hosts^22^. Determining the precise mode of MaRNAV transmission is therefore an important step in disentangling these possibilities. In the absence of a robust *in vitro* system, an investigation of the genetic relatedness of *P. vivax* sub-populations that harbour this virus could be used to further understand both *Plasmodium* and viral transmission dynamics between mosquitoes and humans.

The absence of MaRNAV-1 in *P. cynomolgi* and *P. knowlesi* may reflect distinct evolutionary trajectories between these Asian primate parasites and human *P. vivax*. Phylogenetic analyses position *P. vivax* either as a sister taxon to, or basal to, the Asian primate parasite clade^23^, with recent genomic evidence demonstrating that extant human *P. vivax* represents a bottlenecked lineage that emerged from more diverse African ape parasites^24^. The specificity of MaRNAV-1 to *P. vivax* could reflect either acquisition of this viral association after the divergence from Asian primate *Plasmodium* lineages, or differential retention/loss dynamics during independent host adaptations. African ape *P. vivax*-like parasites exhibit approximately 10-fold greater genetic diversity than human *P. vivax*, consistent with a severe population bottleneck during the establishment of the human parasite lineage^24^.

In contrast, the absence of MaRNAV-1 infection of the other human-infecting *Plasmodium* species that relapse from latent hepatic hypnozoite parasite stages, *P. ovale* and *P. cynomolgi*, suggests that MaRNAV-1 may not be essential in enabling relapse from hypnozoites in *Plasmodium spp*. Nevertheless, the role of MaRNAV-1 in the biology, pathogenesis and transmission of *P. vivax* in humans remains unclear.

Together, our findings expand our understanding of the Matryoshka RNA virus lineage while raising new questions about the virus-parasite relationship. These questions, such as why *P. vivax* is the only known *Plasmodium* species to harbour MaRNAVs and its role in *P. vivax* pathobiology and transmission in humans, have broader implications for both evolutionary biology and public health. Resolving them through expanded sampling and experimental analyses will become increasingly pertinent if global strategies for the elimination of *P. vivax* falter.

## METHODS

### Collection of primary isolates

Whole blood was collected from patients presenting with *P. vivax* malaria in Sabah, Malaysia (n=6)^25^, Papua, Indonesia^26^, including malaria imported to Darwin from this region after 2008^27^ (n=18), Solomon Islands (n=2)^28^ and India (n=1)^29^; *P. falciparum* detected in Sabah (n=5)^30^, Tanzania (n=2)^31^, and imported from Papua, Indonesia (n=2), Mali (n=1), Ghana (n=2), and Cambodia (n=1)^27^; *P. ovale* detected in Sabah, Malaysia (n=1)^30^ and Papua, Indonesia (n=6)^32^; and *P. malariae* detected in Sabah, Malaysia (n=2)^33^ and Papua, Indonesia (n=6)^32^. Ethics approval was obtained for blood collections from patients with malaria as listed in each of the studies cited above. We also tested a laboratory-adapted strain of *P. cynomolgi Berok*^*34*^ for the presence of RNA viruses including MaRNAVs.

### Total RNA sequencing of primary samples

Total RNA was extracted from the primary human blood samples using the RNeasy Universal kit (QIAGEN). Subsequent gDNA removal was performed on total RNAs using DNAse I treatment (QIAGEN). Pooled RNA was prepared for total RNA sequencing with the TruSeq stranded library (Illumina) and Globin-Zero gold kit (Illumina). Libraries were sequenced as paired-end 150 bp using either the Illumina NovaSeq 6000 S4 or SP lane (Australian Genome Research Facility, Melbourne). PCR was used to confirm the presence of MaRNAV-1 in individual primary samples (**Fig. S1**).

### Bioinformatic processing

#### SRA Screening

All publicly available *Plasmodium* RNA SRA libraries generated through RNA-Seq were downloaded from NCBI according to the following criteria: (i) not included in the original Charon, et al., study^1^; (ii) at least 0.4GB in size. These libraries were then screened for the presence of MaRNAV-1-7. The *P. falciparum* SRA libraries were downloaded using SRA Toolkit v3.0.3.

#### Trimming and assembly

Primary libraries were trimmed using TRIMMOMATIC v0.38^35^ and TruSeq3 paired-end adapters (minimum length = 25). Libraries from the SRA were trimmed using Cutadapt^36^ v1.8.3 (-u 5 -U 5 -q 25 -m 25). *Plasmodium* reads were removed from primary libraries using Bowtie2^37^ (**Table 3**). All trimmed and filtered reads were assembled using MEGAHIT^38^ v1.2.9 with default parameters.

**Table 3:** Reference genomes used to remove *Plasmodium* reads from primary libraries.

| Host | Reference genome |
| --- | --- |
| *P. ovale* | GCA_900090025.2_PowCR01 |
| *P. malariae* | GCF_900090045.1_PmUG01 |
| *P. falciparum* | GCF_000002765.4_ASM276v2 |
| *P. vivax* | Pv_GCF_000002415.2_ASM241v2 |
| *P. knowlesi* | GCF_000006355.1 |

### Targeted virus discovery

We used two approaches to search for MaRNAV-like contigs in the libraries. For and assembled them using MEGAHIT v1.2.9 with default parameters.

For the libraries generated from the primary samples and the *P. vivax* SRA, we created a custom DIAMOND database comprising all known MaRNAVs (both RdRp and hypothetical protein segments) and screened our libraries against this database using DIAMOND v2.1.6^14^. All contigs that returned a significant hit to this database (either e-value < 0.001 and default sensitivity for *P. falciparum* SRA libraries or e-value < 1e-5, --ultra-sensitive for all others) were checked against the NCBI non-redundant (nr) protein database (as of March 2025) again using DIAMOND BLASTx, and false positives (i.e., contigs with detectable sequence similarity to cellular genes, e-value < 1e-5, --very-sensitive) were removed.

### RT-qPCR confirmation of MaRNAV-1 in primary isolates

The primary isolates were pooled for total RNA sequencing. We therefore used RT-qPCR as reported previously^1^ to test for the presence of MaRNAV-1 in individual isolates that comprised pools in which MaRNAV-1 was detected with RNASeq. Briefly, cDNA was synthesised using the SuperScript IV VILO reverse transcriptase (Invitrogen) and amplification was performed using virus-specific primers^1^ and the Platinum SuperFi DNA polymerase (Invitrogen) (35 cycles, annealing temperature 64C). The human gene RSP18 was used as the positive control.

### Assessment of host composition

We analysed the composition of *P. falciparum* libraries in which we had detected the MaRNAV-1 RdRp segment using Kraken2 v2.1.3^39^. Read counts for Opisthokonta (i.e., animals and fungi), *Plasmodium (Laverania), Plasmodium vivax*, and remaining cellular organisms were extracted from the report file in R. The ratio of *P. vivax* to *P. falciparum* reads was calculated using these read counts.

### MaRNAV-1 frequency estimates

#### Percent positivity

To estimate the frequency of association between MaRNAV-1 and *P. vivax*, we filtered the complete data set of *Plasmodium* libraries that we screened to exclude all samples except those known to have been collected from human infections. This data set was further filtered to remove instances of repeated sampling from the same individual for longitudinal study, retaining samples taken prior to antimalarial treatment where possible.

We first tabulated the samples in this refined data set by country. We then measured frequency as the number of libraries (SRA) or samples (primary) with MaRNAV-1 RdRp per sample count per country and reported this figure as a percentage.

#### Binomial probability function

To measure the probability of observing 120 or fewer MaRNAV-1-positive samples among the 1453 human-infecting *P. falciparum* libraries screened here (8.26% frequency), we used the binomial distribution function implemented in R (pbinom, **Eq. 1**). We assumed a true frequency of 20.89%, as this was the weighted average in our data set.

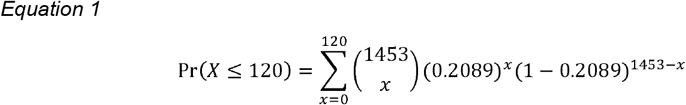

### Translation and ORF identification

Nucleotide sequences of the MaRNAV-1 RdRp genes identified in this study were translated using the EMBOSS getorf tool (https://www.bioinformatics.nl/cgi-bin/emboss/getorf). Open reading frame (ORFs) detection was enabled between stop codons and were limited to a minimum length of 300bps. For contigs that returned multiple putative ORFs that met these parameters, the longest ORF was selected. To assess the accuracy of this selection, a preliminary multiple sequence alignment was performed using MAFFT^40^ v.7.490 with default parameters implemented in Geneious Prime. Incorrect ORFs were identified, removed, and replaced manually.

### Phylogenetic analysis

#### Phylogenetic position of MaRNAVs within the Narnaviridae

An initial reference data set was compiled by screening the RdRp segment of MaRNAV-1 against the NCBI nr database and downloading the first 100 hits. The amino acid sequences of all published MaRNAVs^1-4^ were also downloaded. Sequences were aligned with MAFFT^40^ v7.490 using default parameters and implemented in Geneious Prime. Ambiguous sites were removed using trimAl^41^ v1.4.1 (‘gappyout’). Phylogenetic analysis (**Fig. S2**) was performed using the maximum likelihood method implemented in IQ-TREE^42^ v1.6.12 with the ModelFinder limited to LG models with 1000 ultrafast bootstraps and SH-aLRT test utilised to measure support. LG+RF+R7 was selected as the best-fit model according to the Bayesian Information Criterion (BIC). The resulting phylogenetic tree was rooted on the non-MaRNAV clade.

#### Phylogeographic mapping

A data set of all MaRNAV-1 RdRp sequences was compiled (n=1,110; one of which was identified previously^1^). We again used MAFFT, trimAl, and IQ-TREE to infer a phylogenetic tree, this time retaining MaRNAVs-2, -3, -6, and -7 as the outgroup. All parameters for these methods were retained except that we did not restrict the ModelFinder. LG+R4 was selected as the best-fit model according to BIC. The geographic (i.e., country of sampling) of each MaRNAV was then mapped onto the tips of the phylogenetic tree.

To visualise the phylogeny, we used the R libraries ape, ggtree, and gheatmap implemented in Rv4.4.0. We assigned heatmap colours according to the designated country (if known) and microbial host obtained from our primary sample or SRA metadata. We refined the visualisation with Adobe Illustrator.

#### Root-to-tip analysis of evolutionary rates

To estimate the evolutionary rate of MaRNAV-1 in this data set, we retained only MaRNAV-1 RdRp sequences for which at least the year of sampling was known (n=794). When only the year was known, we assigned June 1 of that year as the sampling date. All sequences were aligned with MAFFT v7.490 using default parameters (final alignment: 6,564 nucleotides in length, 93.2% pairwise identity). We inferred a maximum likelihood phylogenetic tree using IQ-TREE v1.6.12 (1000 ultrafast bootstraps and sh-aLRT). ModelFinder was unrestricted and selected TPM2u+F+R5 according to the BIC score. Evolutionary rate (i.e. the number of nucleotide changes per site per year) was estimated by plotting root-to-tip genetic distances against the year of sampling with TempEst v1.5.3^43^, employing the heuristic residual mean squared function to find the best-fitting root (**Fig S3**).

## ACKNOWLEDGMENTS

This work was funded by a National Health & Medical Research Council (NHMRC) Investigator grant (GNT2017197) to E.C.H., an Australian Research Council (ARC) Discovery Project grant (DP240101313) to E.C.H. and M.E.P. We thank Ethan Mandojana for assistance with data cleaning and our colleagues at the National Institute of Health Research and Development and Royal Darwin Hospital for assistance with the studies enrolling patients with malaria.

## AUTHOR CONTRIBUTIONS

M.E.P., J.C., M.G., J.W., E.C.H., and N. M. A. conceived this study. M.G., J.W., B.R., R.P., T.W., E.K., J.M., and B.E.B. collected the primary samples. M.E.P., J.C., R.H.P., and K.S. performed the analyses. M.E.P., E.C.H., and N.M.A. drafted the manuscript. All authors reviewed and edited the manuscript.

## DATA AVAILABILITY

The MaRNAV-1 sequences identified in this study are available on GenBank (accs: XXXX-XXXX).

