## Supplementary material for "Mapping the global distribution and spread of the *Plasmodium vivax*-associated virus MaRNAV-1": Fig S1

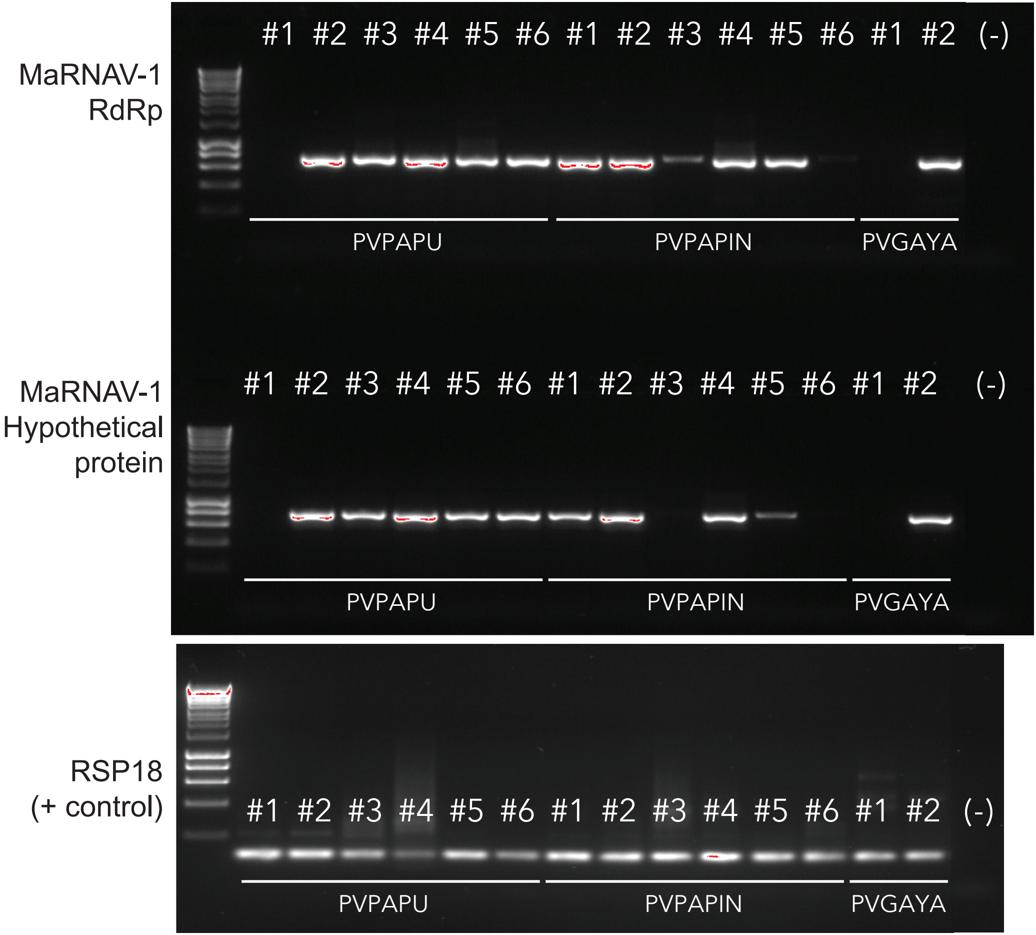


**Figure S1 Visualisation of MaRNAV-1 segments in primary human blood samples with *P. vivax* infection that were pooled for sequencing.** Abbreviations: PV = *P. vivax*; PAPU, PAPIN = Papua, Indonesia; GAYA = Gaya Island, Sabah, Malaysia
