## Supplementary material for "Mapping the global distribution and spread of the *Plasmodium vivax*-associated virus MaRNAV-1": Fig S2

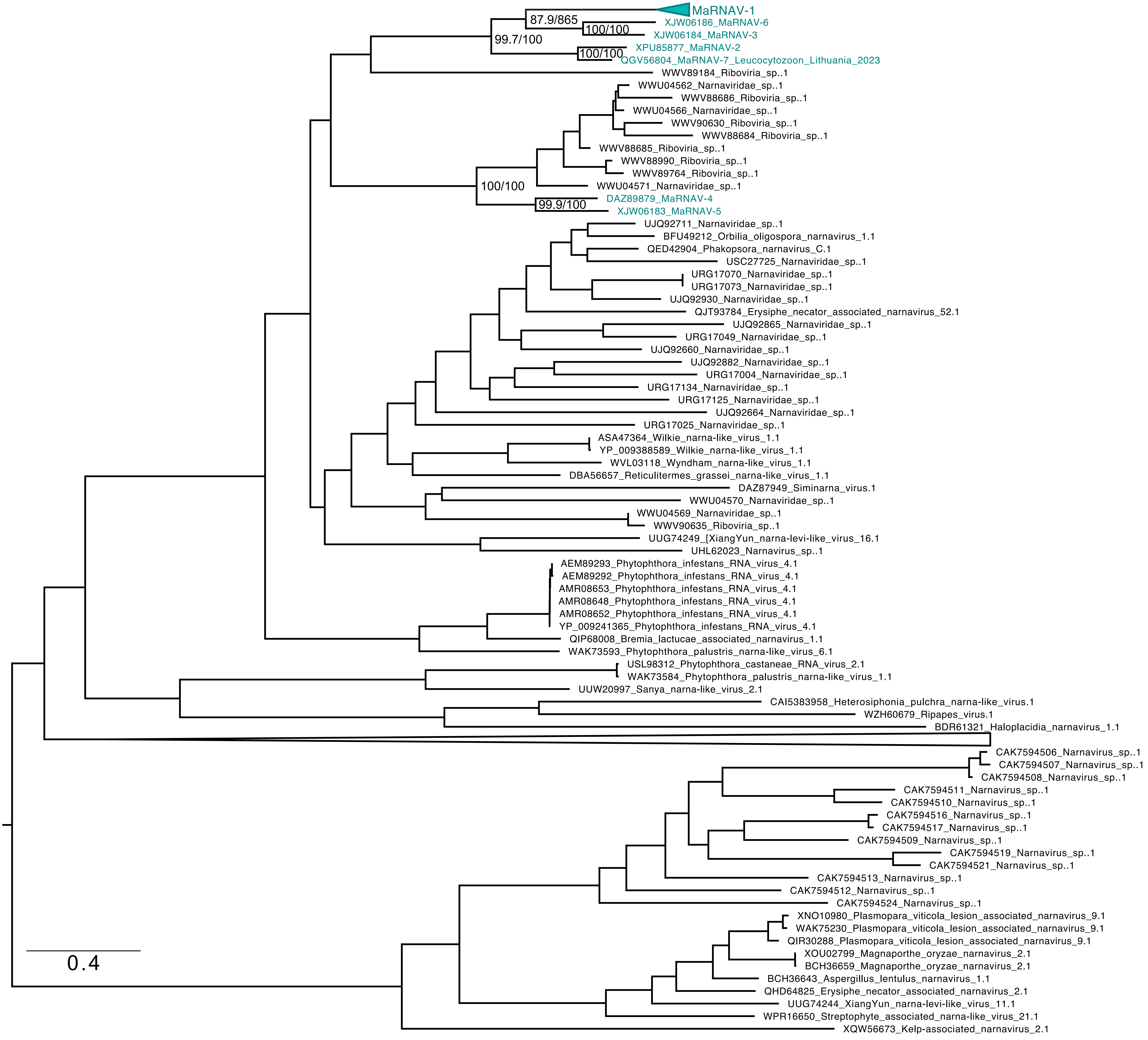


**Figure S2 Relationship of known MaRNAVs within the *Narnaviridae*.** Mid-point rooted maximum likelihood phylogeny scaled by the number of amino acid substitutions. MaRNAV lineages are denoted by green tips. Support values are shown at select nodes as sh-aLRT/UFBoot.
