## Supplementary material for "Mapping the global distribution and spread of the *Plasmodium vivax*-associated virus MaRNAV-1": Fig S3

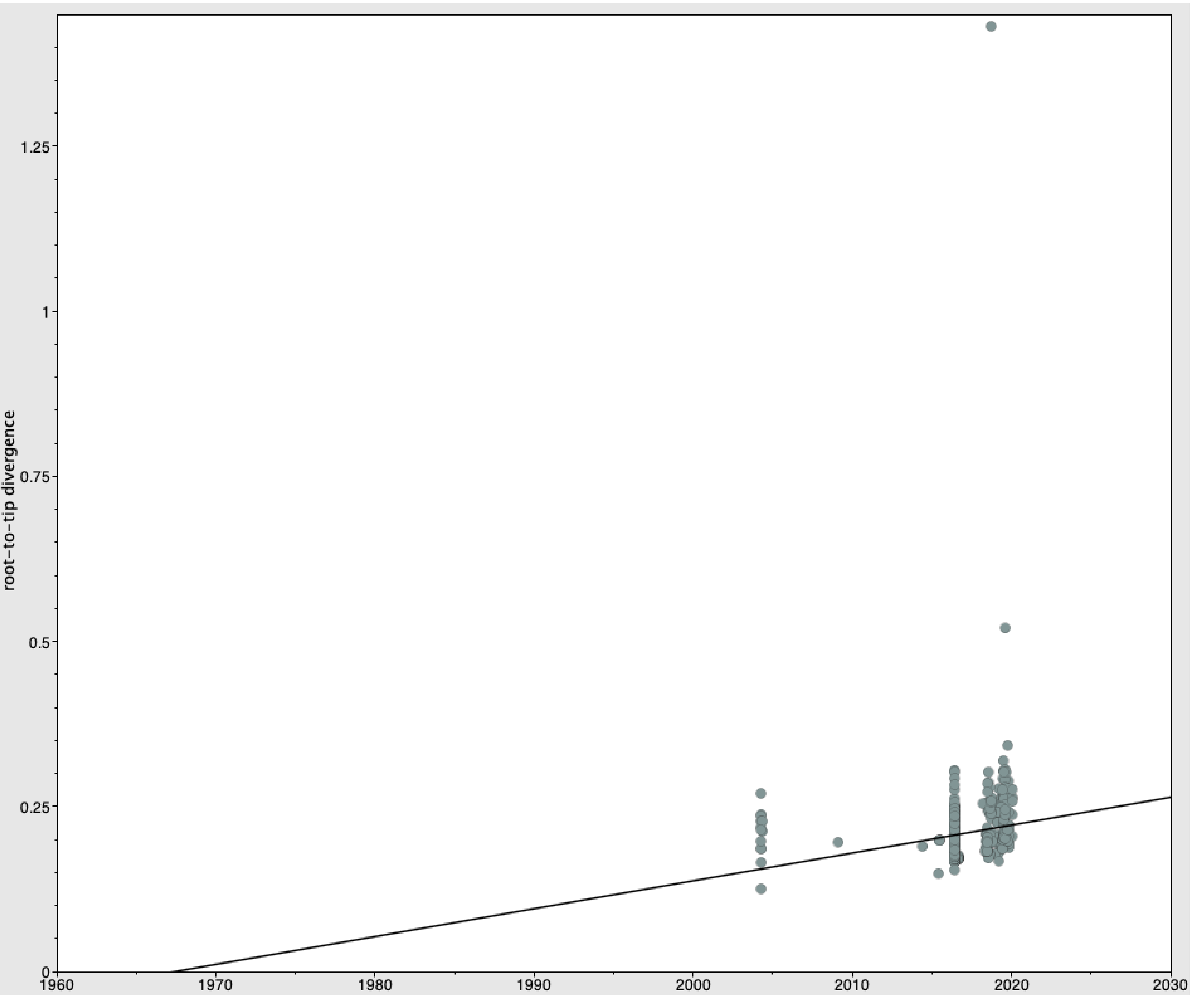


**Figure S3 Root-to-tip divergence of MaRNAV-1.** The best-fitting root was determined using the heuristic residual mean squared function. The results were visualised in TempEst. Year is shown on the Y axis..
